# *In situ* structural analysis of mammalian cells using a 200 kV electron cryomicroscope – implications for research infrastructure

**DOI:** 10.1101/2024.12.06.627167

**Authors:** Piotr Szwedziak

## Abstract

**Background:** Electron cryotomography is a powerful imaging technique allowing for studying functional cellular modules in their native environment with macromolecular resolution. However, it requires access to complex and expensive instrumentation, typically a 300 kV electron cryomicroscope equipped with an energy filter. Simpler and cheaper 200 and 100 kV instruments have been successfully used for single particle cryoEM analyses, which has helped to democratize the technique and broaden access. It has not been systematically studied if 200 kV electron cryomicroscopes can deliver meaningful and interpretable data with respect to electron cryotomography applications.

**Methods:** Here, we set out to investigate if a 200 kV electron cryomicroscope without an energy filter can be utilized for *in situ* structural studies of mammalian cells by electron cryotomography of thin cell edges followed by extensive image analysis including segmentations, subtomogram averaging and molecular sociology studies of lipid droplets.

**Results:** We demonstrate that the resulting tomograms of thin edges of U2OS cells are of sufficient quality to annotate the contents of the cell and observe spatial inter-relationships among macromolecules. In particular, we undertook a molecular sociology analysis of lipid droplets and addressed their subcellular distribution and interactions with other organelles. Additionally, we performed subtomogram averaging of purified 70S ribosomes that resulted in ∼15 Å resolution 3D reconstruction. Finally, we examined geographical distribution and scientific output of the two most common electron cryomicroscopy platforms and deduced that 200 kV instruments are heavily underutilized with respect to electron cryotomography applications.

**Discussion:** This study demonstrates that 200 kV electron cryomicroscopes can be utilized for structural cell biology studies by electron cryotomography. Given the favorable ratio of their versatility versus costs we foresee that 200 kV electron cryomicroscopes will become workhorses of local electron cryomicroscopy facilities.

## Introduction

It has been a decade since the resolution revolution in the electron cryomicroscopy (cryoEM) field occurred (Kühlbrandt, 2014). Single particle analysis (SPA) cryoEM has been a tremendously successful structural biology technique (as evidenced by the exponentially growing number of relevant entries in the Electron Microscopy Data Bank) enabling routine determination of high-resolution 3D structures of purified macromolecules such as heterogenous or flexible protein complexes (Punjani & Fleet, 2023; Schwab et al., 2024).

However, biological functions are not executed by individual molecules in isolation but rather complex networks of interactions between all the cellular components, including proteins, nucleic acids and lipids, are responsible for maintaining cellular homeostasis. Therefore, the need for structural studies by cryoEM in unperturbed cellular environments is steadily growing (Nogales & Mahamid, 2024). Electron cryotomography (cryoET) is a cryoEM modality that provides 3D molecular resolution images of isolated/*in vitro* reconstituted cellular components as well as intact cellular environments, which is known as *in situ* cryoET. CryoET preserves the full spectrum of conformations and interactions of each molecule, enabling detailed ultrastructural analyses of cells and tissues in health and disease that has been described as ‘visual proteomics’ (Beck & Baumeister, 2016; Förster et al., 2010). If structures of interest are present in multiple copies within a tomogram, they can be extracted, aligned and averaged in a process called subtomogram averaging, which is analogous to SPA but makes use of 3D volumes rather than 2D projections. In favorable conditions this can yield a near-atomic resolution reconstructions (Tegunov et al., 2021). Additionally, the position and context of molecules and how they relate to the cellular context are available.

Given that the mean free path for inelastic scattering for 300 kV electrons is ∼300-400 nm (Martynowycz et al., 2021; Rice et al., 2018), a critical requirement and limitation for cellular cryoET is sample thickness. The resulting tilt series images must have enough signal to noise ratio in order to be reliably aligned when reconstructing the 3D volume (tomogram). Therefore, initially the analyses were limited to *in vitro* specimens, bacteria or thin edges of eukaryotic cells (Medalia et al., 2002; Szwedziak et al., 2014; Szwedziak & Pilhofer, 2019). In the last ten years techniques based on dual-beam FIB-SEM instruments have become available to prepare thin lamellae of cells and tissues for subsequent cryoET analyses (Rigort & Plitzko, 2015).

Complex sample preparation techniques, fragmented data processing pipelines and lower throughput, as compared to SPA cryoEM, are primary reasons why cryoET still lags SPA in terms of maturity and output. Cellular cryoET is a costly technique as it frequently requires access to high-end 300 kV TEMs equipped with energy filters that remove inelastically scattered electrons in order to increase the signal-to-noise ratio. In the future the costs will become even more prohibitive given the implementation of laser phase plates and Cc correctors that will yield an ultimate, cryoET purpose-built instrument (Russo et al., 2022).

Given that the mean free path at 200 kV is ∼260 nm and inspired by the study (Martynowycz et al., 2021) suggesting that useful cryoEM data can be collected up to twice the inelastic mean free path without the need for energy filters, we set out to investigate if a 200 kV TEM equipped with a direct electron detector is sufficient to provide meaningful cryoET data.

Here, we imaged by cryoET thin edges of mammalian U2OS cells using a 200 kV Glacios G1 cryoTEM fitted with a Falcon 3EC camera without an energy filter. The tomograms enabled us to observe a plethora of cellular structures ranging from mitochondria through cytoskeletal filaments to macromolecular complexes, e.g. ribosomes. By combining segmentation with subtomogram averaging we generated detailed ultrastructural cellular snapshots that might be useful in addressing various cell biology problems, e.g. lipid droplets (LDs) metabolism. LDs are central to lipid and energy homeostasis and are important storage organelles (Olzmann & Carvalho, 2019). LDs are known to originate in the endoplasmic reticulum (ER) and associate with other cellular structures and a myriad of proteins (Olzmann & Carvalho, 2019). To better understand the role of LDs in cellular metabolism we undertook a molecular sociology analysis of LDs and addressed their subcellular distribution and interactions with other organelles. We also provide visual evidence for direct interactions between LDs and membrane trafficking proteins. Finally, we obtained a ∼15 Å subtomogram average of purified and vitrified *Escherichia coli* 70S ribosomes.

The current trend towards democratizing cryoEM in the SPA field is gaining momentum and it has been demonstrated that 200 kV (Herzik et al., 2017) and 100 kV (McMullan et al., 2023) cryoTEMs are sufficient to obtain high-quality SPA cryoEM data. Here, we prove that 200 kV TEMs without an energy filter can be utilized for cellular cryoET investigations. We suggest that 200 kV cryoTEMs can be cost-effective workhorse cryoEM instruments capable of providing complementary information from both major cryoEM modalities and with the recent development of electron detectors optimized for lower electron energies (Chan et al., 2024; McMullan et al., 2023) their performance and significance will steadily grow.

## Results

### CryoET imaging of mammalian cells

CryoET has been frequently employed to address cell biology problems that otherwise were not accessible to other imaging techniques, e.g. coronavirus replication (Wolff et al., 2020) or detailed membrane architecture (Wozny et al., 2023). To ensure optimal performance and data quality these studies made use of complex 300 kV TEMs equipped with energy filters. Due to prohibitive costs of purchase and maintenance, such instruments are predominantly available in major research centers with limited access. Given that the mean free path for inelastic scattering at 200 kV is ∼260 nm, which is comparable to the average cryoET specimen thickness, we set out to investigate if a 200 kV TEM without an energy filter can deliver interpretable cryoET results. Specifically, we were interested in understanding the interplay and spatial arrangement of LDs in the context of intact cells.

U2OS cells were seeded on electron microscopy grids with a hole carbon support film and vitrified by plunge-freezing. These cells are known to adopt a flat, pancake-like shape that offers a significant cryoET imaging surface over thin cells areas without the need for micromachining by cryoFIB. The grids were imaged by a Glacios G1 cryoTEM equipped with a Falcon 3EC camera without an energy filter. The cells could be easily recognized at low magnifications (**Fig. 1a, left**) with the cell nucleus surrounded by the electron transparent cytoplasmic regions. This allowed for a relatively straightforward setup of automated tilt series acquisition and great care was taken to tune the 2-condenser lens system to parallel illumination conditions. It was possible to acquire multiple tilt series from a single cell, and the final thickness of the 3D volume was dictated by the local cell thickness (**Fig. 1a, small panels**). The imaged areas were structurally well preserved and showed no sign of crystalline ice contamination. In total, 64 cells were imaged over which we recorded almost 230 tomograms with an average thickness of 240 ± 75 nm (**Fig. 1b**). Most tomograms were between 200 and 250 nm thick, with the thinnest being 50 nm and thickest 400 nm (**Fig. 1c**). Attempts to record tilt series on thicker areas failed due to the low contrast of resulting images and poor tracking. To assess the quality of the data quantitively we fitted the contrast transfer function (CTF) on the 2D zero tilt images and using CTFFIND4 we estimated the resolution up to which the CTF is fitted reliably. We found that the precision of the fit is linearly dependent on the local cell thickness (**Fig. 1d**). Encouraged by these results we moved on to a more detailed inspection of the tomograms.

**Figure 1.**
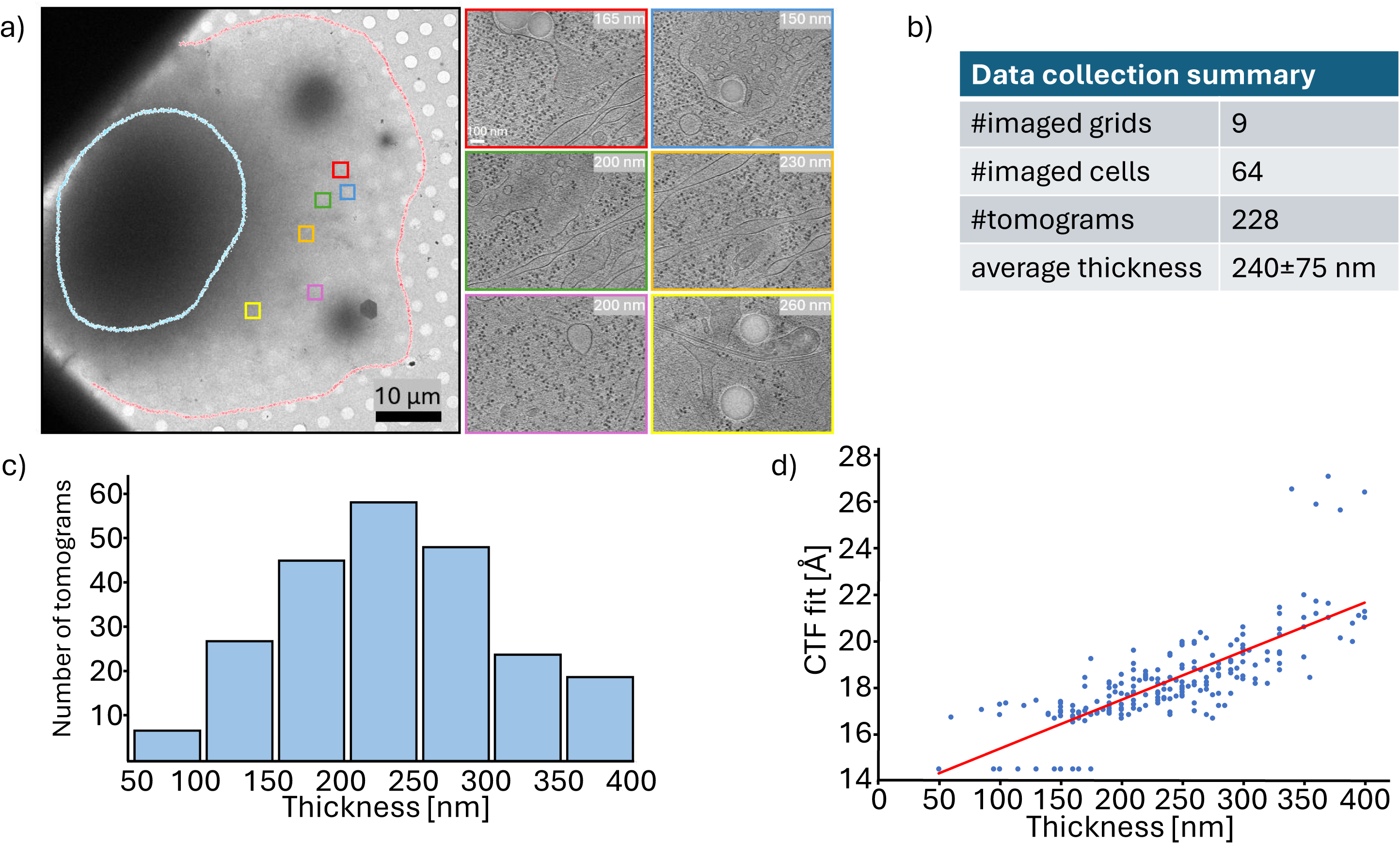
CryoET analysis of intact mammalian cells using a 200 kV cryoTEM without an energy filter. a) A low magnification overview of a U2OS cell seeded on a Quantifoil grid and vitrified in liquid ethane. The electron opaque area encircled in blue corresponds to the nucleus/perinuclear area. The cell boundaries are encircled in red. Six tilt series were acquired in the electron-transparent area corresponding to the thin cell peripheries. Central slices of reconstructed tomograms from each area are shown in the panels on the right and color-coded to mark the acquisition areas. Measured tomogram thicknesses are displayed. Scale bars: 10 µm and 100 nm. b) Table summarizing the dataset acquired in this study. c) Histogram of the tomogram thicknesses. The thickness varied from 50 to 400 nm with the most populated class in the range of 200-250 nm. d) Plot of resolution up to which CTF was reliably fitted for a central tilt in every tilt series against the tomogram/local cell thickness. The red line shows a linear fit through all data points.

### Molecular atlases of native cellular environments

Since the aim of this study was to generate low nanometer resolution 3D volumes of intact mammalian cells in order to qualitatively and quantitatively describe LDs molecular neighborhoods, we adopted an appropriate data collection strategy (**Table 1**): we utilized relatively high defocus (range from −5 to −7 μm) and high total dose (140 eÅ^-2^) to boost image contrast and large pixel size (5.1 Å) to benefit from the broad cellular context provided by the large field of view (∼2 μm). The figures presented were prepared using tomograms generated by SIRT (Simultaneous Iterative Reconstruction Technique) as it offers high contrast that is useful for downstream cellular tomography applications such as segmentations and annotations. The quality of the tomograms can be judged from the clearly distinguishable cellular features, e.g.: microtubules, intermediate filaments, actin filaments, lipid tubules, ribosomes (**Fig. 2a, left panels**). A representative tomogram of thickness of 180 nm is shown in **Fig. 2a (left large panel)** and **Movie S1**. In the total volume of ∼0.72 μm^3^ we could find the three classes of intracellular filaments (microtubules, intermediate filaments, actin) intertwined with each other as well as with membranous structures of the endoplasmic reticulum (ER) and ribosome particles. Subsequently we trained a neural network within the EMAN2 software package (Chen et al., 2017) to recognize lipid bilayers, intermediate filaments, double membranes of mitochondria, microtubules and ribosomes and applied it to the tomograms (**Fig. 2a, middle panels**). This allowed for reproducible identification of features of interest and their separation from the background. By applying the neural network, we identified and automatically extracted **∼**1500 ribosomal particles and subjected them to subtomogram averaging. This resulted in a moderate resolution 3D reconstruction at 38 Å with the conservative threshold of 0.5 for FSC (**Fig. 2a, small panel**). Including more particles in the average didn’t improve the resolution suggesting that the dataset is limited by data collection strategy used (high defocus and large total electron dose) and sample thickness. Importantly, resolution is not the best criterion for data quality in cryoET as the contextual information provided by cellular tomograms can be of higher significance. Accordingly, by mapping the averaged structure to the calculated positions and orientations of each particle in the tomograms and combining it with segmentations of other cellular components we generated 3D molecular maps of thin edges of intact mammalian cells (**Fig. 2a, right panel, Movie S2).** Such maps are visually appealing to human perception and easier in interpretation than raw tomograms. We find the prospect of producing detailed annotations of 3D cryoET volumes obtained by imaging with a 200 kV cryoTEM without an energy filter particularly exciting since such segmentations aid interpretation and are essential for many downstream cryoET analysis steps, e.g. observing spatial inter-relationships among macromolecules.

**Figure 2.**
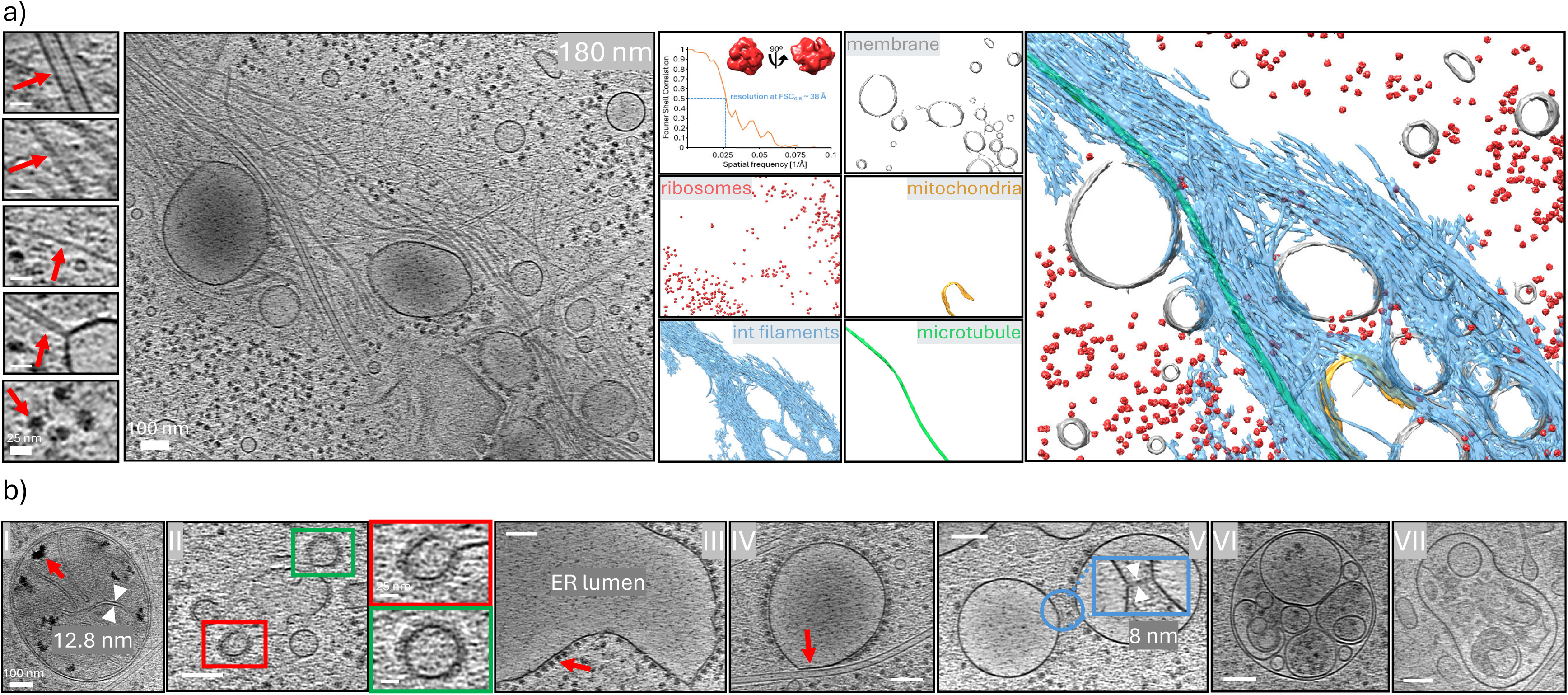
CryoET at 200 kV provides molecular resolution atlases of intact mammalian cells. a) Central slice (14 nm thick) through a tomogram of an intact U2OS cell with a local thickness of 180 nm (left large panel). Small panels on the left display common mammalian cell features that were identified in the tomogram, from top to bottom: microtubules, intermediate filaments, actin, membrane tubules and ribosomes (red arrows). By employing machine learning-based approaches we could identify, extract and average subvolumes containing ribosome particles and map the resulting low nanometer resolution average back in the tomogram (middle panels). The quality of the tomogram was sufficient to segment other components of the cell: membranes, mitochondria, intermediate filaments and microtubules (middle panels). By combining all the segmentations, we generated a simplified model of the cell that can aid interpretation (right panel). Scale bars: small panels – 25 nm, main panel – 100 nm. b) Examples of some of the functional cellular modules that could be readily identified in the tomograms: I) Mitochondrion with calcium phosphate inclusions (red arrow). Measured cristae diameter is highlighted in white. II) Putative COPI-coated vesicles budding off an early endosome. Color-coded are enlarged views of highlighted areas at a different tomographic slice. COPI coat is characterized by a very dense and compact appearance. III) endoplasmic reticulum (ER) with its empty lumen and the cytoplasmic surface covered with ribosomes (red arrow). IV) ribosome-associated vesicle (RAV) covered densely with ribosome particles except where it interacts with a microtubule filament that ultimately leads to a local ribosome deprivation and membrane flattening of otherwise spherical structure (red arrow). V) RAVs and ER subcompartments are interconnected by a dense network of thin membrane tubules that are 8 nm in diameter, as highlighted in the enlarged view that is a snapshot at a different tomographic slice of the area encircled in blue. VI) A membrane-embedded compartment that could be an autophagosome. VII) A bell-like shaped membrane compartment with internal debris that could be an endolysosome that is a result of the fusion of a lysosome with a multivesicular body. Scale bars: major panels – 100 nm, enlarged views – 25 nm.

**Table 1.** CryoET data collection.

|  |  |
| --- | --- |
| Microscope | Thermo Fisher Glacios G1 |
| Camera | Falcon 3EC |
| Energy filter | N/A |
| Magnification | 28000 |
| Voltage (kV) | 200 |
| Electron exposure (eÅ<sup>-2</sup>) | 140 |
| Defocus range (μm) | 5.0-7.0 |
| Pixel size (Å) | 5.1 |
| Tilt range | ±54° |
| Increment | 3° |

We went on to evaluate if the 200 kV cryoET data allow for reliable identification of other known subcellular structures (**Fig. 2b**). In 13 tomograms we recognized mitochondria that adopted either spherical or rod-shaped morphologies (**Fig. 2b (I), Movie S3 and Fig. S1**). The measured cristae width was 12.8 nm, which stays in accordance with the previously published 14.5±5.8 nm (Fry et al., 2024). Within the mitochondrial matrix we could observe abundant granules characterized by high contrast – we attributed those densities to calcium phosphate storage, as first described in (Wolf et al., 2017). Calcium phosphate granules are used to regulate calcium homeostasis and to maintain mitochondrial function. Similar densities were observed by cryoET in neuronal mitochondria (G. H. Wu et al., 2023). CryoET at 200 kV could be an excellent diagnostic platform for *in situ* organelle phenotyping as it has been shown that mitochondrial ultrastructure, including its membrane architecture, has profound implications for human disease (Navaratnarajah et al., 2021).

CryoET at 200 kV was sufficient to resolve coated vesicles/buds that play a role in transport between various cellular compartments (**Fig. 2b (II), Movie S4**). The overall morphology and the localization in the cell peripheries indicate that the presented membrane-bound compartment could be an early endosome. A closer look at the dense protein coat and comparison to the previously published cryoET data (Bykov et al., 2017) suggests it is COPI. COPI coat is known to play an important role in mediating transport between the Golgi and ER within the secretory pathway but it has also been demonstrated to act in early endosome function (Daro et al., 1997). The observed buds/vesicles had an average diameter of 55±7 nm (n=7), which stays in accordance with other reports (Bykov et al., 2017; Faini et al., 2012). To fully appreciate the complex membrane architecture, we recommend seeing **Movie S4**.

In the imaged cells we found extensive areas of ER. The observed ER is characterized by the lumen deprived of other intracellular organelles/macromolecular complexes and is tightly coated on the cytoplasmic face by ribosomes where individual particles could be discerned (**Fig. 2b (III) and Movie S5**). The ER network consisted of cisterna interconnected by an array of branching tubules. In 10 tomograms we also identified a recently reported form of ER – ribosome associated vesicles (RAVs). RAVs were originally described as dynamic ER subcompartments found primarily in the periphery of secretory cells (Carter et al., 2020). Here, we found numerous RAVs interacting tightly with microtubules that in the most severe cases triggered membrane flattening (**Fig. 2b (IV) and Movie S6 and Fig. S2**). RAV-bound ribosomes were excluded from the RAV-microtubule interface. The observed RAVs were interconnected via membrane tubules that were 8 nm in diameter (**Fig. 2b (V) and Movie S7**). Membrane-bound polyribosome, which is a cluster of ribosomes bound to an mRNA, was found on the RAV surface (**Fig. S3 and Movie S8**). Given that actively translating mRNAs are frequently associated with multiple ribosomes we suspect that RAVs might be sites of membrane/extracellular protein synthesis, similarly to the canonical ER.

We also observed multilamellar compartments containing vesicles as well as membrane sheets and other cellular debris/dense deposits. These are most likely degradative compartments, for example autophagosomes (**Fig. 2b (VI) and Movie S9**) or endolysosomes (**Fig. 2b (VII) and Movie S10**), as described in (Foster et al., 2022). The luminal density within these organelles was highly variable, suggesting that their content is different and not well defined.

### Detailed analysis of LDs

In 84 tomograms we noticed the presence of LDs. They are characterized by an amorphous, electron-dense core made of neutral lipids with a single phospholipid layer on the outside, which is manifested as the electron-dense line delineating the LDs circumference in the tomograms (**Fig. 3a, top panel and Movie S11**). In total we recognized 231 LDs that were 205±70 nm in diameter. LDs were often found in clusters (**Fig. 3a, bottom panels and Movie S12**) and 29% of LDs stayed in direct contact with each other, which frequently resulted in shape changes and flattening (**Fig. 3a, bottom right panel and Fig. S4**), as reported previously (Ganeva et al., 2023). ER double leaflet is the site of LDs synthesis (Fujimoto & Parton, 2011) and they remain tightly associated with the ER throughout their life cycle (**Fig. 3b, top right panel and Movie S13**): indeed, 69% of the cytoplasmic LDs were in contact with ER whose membrane curvature followed LD’s shape (**Fig. 3b, top left panel, Movie S14**). Interestingly, 5.6% of the observed LDs were positioned in the ER lumen (**Fig. 3b, top right panel and Fig. S5**). The luminal LDs are the least characterized members of the mammalian LD family, and more investigations are needed to establish what triggers LDs movement towards either cytoplasmic or luminal side. LDs association with ER was described as very promiscuous (Valm et al., 2017) and we could observe several distinct instances of this interaction. For example, luminal and cytoplasmic LDs in direct contact presumably during material exchange (**Fig. 3b, bottom middle panel**) or cytoplasmic LDs impacting the ER membrane and remaining at a distance of 10-12 nm that is sufficient to accommodate proteins mediating the interaction (**Fig. 3b, bottom right panel**). The lipid monolayer is a busy hub for interaction with proteins that frequently are LD-specific (Olarte et al., 2022) and mediate LDs interactions with various organelles (Olzmann & Carvalho, 2019): for example, 3% of all LDs interacted with mitochondria (**Fig. 3c, left panel and Movie S15**) and this association has been reported before as being crucial for metabolism (Fan & Tan, 2024). 40% of the identified LDs interacted with microtubules (**Fig. 3c, middle bottom panel, Movie S16 and Fig. S6**). Direct physical contact between LDs and microtubules can aid with transport of LDs and facilitate finding their cognate organelles. Rapid LD movement on microtubules has been observed by time-lapse fluorescent light microscopy imaging (Gross et al., 2000; Kilwein & Welte, 2019). A small fraction (1.7% of total observed LDs) of LDs interacted with putative lysosomes and MVBs (**Fig. 3c, middle top and right panels**). These LDs are most likely destined for degradation by lysosomal acid lipase and the resulting fatty acids can enter metabolic pathways in mitochondria (Zhang et al., 2022). We could discern direct association of membrane trafficking proteins on the LDs surface. COPI has been demonstrated to act directly on lipid droplets *in vitro* (Thiam et al., 2013) and *in vivo* to enable their connection to the ER for protein targeting (Wilfling et al., 2014). Here, we observed the dense protein coating on the surface of an LD that promoted budding of lipid monolayer resulting in an empty void between the LD core and the monolayer (**Fig. 3d left panels and Movie S17**). Based on the protein coat morphological similarity to previously published coated vesicles (Bykov et al., 2017) we assume that the observed protein density might be attributed to COPI. It is likely that COPI-induced membrane deformation could help establish continuous membrane bridges between the LD and outer ER membrane leaflet (Wilfling et al., 2013), which has been observed in our tomograms (**Fig. 3b, bottom left panel**). Here, the LD is tightly embedded within the lipid bilayer. Another kind of vesicle protein coat, clathrin, has been detected in LDs by mass spectrometry (Bartz et al., 2007; Nonoyama et al., 2019) and our study provides visual confirmation of this interaction (**Fig. 3d, middle and right panels**). The triskelion building block assembling into pentagonal and hexagonal lattice was readily visible and the overall arrangement was similar to clathrin cage architecture reported by SPA cryoEM (Fotin et al., 2006). The role of clathrin in association with LDs is unclear but it has been suggested that it might be important for lipid hydrolysis as it facilitates lysosome formation that is essential for lipid droplet digestion (Nonoyama et al., 2019).

**Figure 3.**
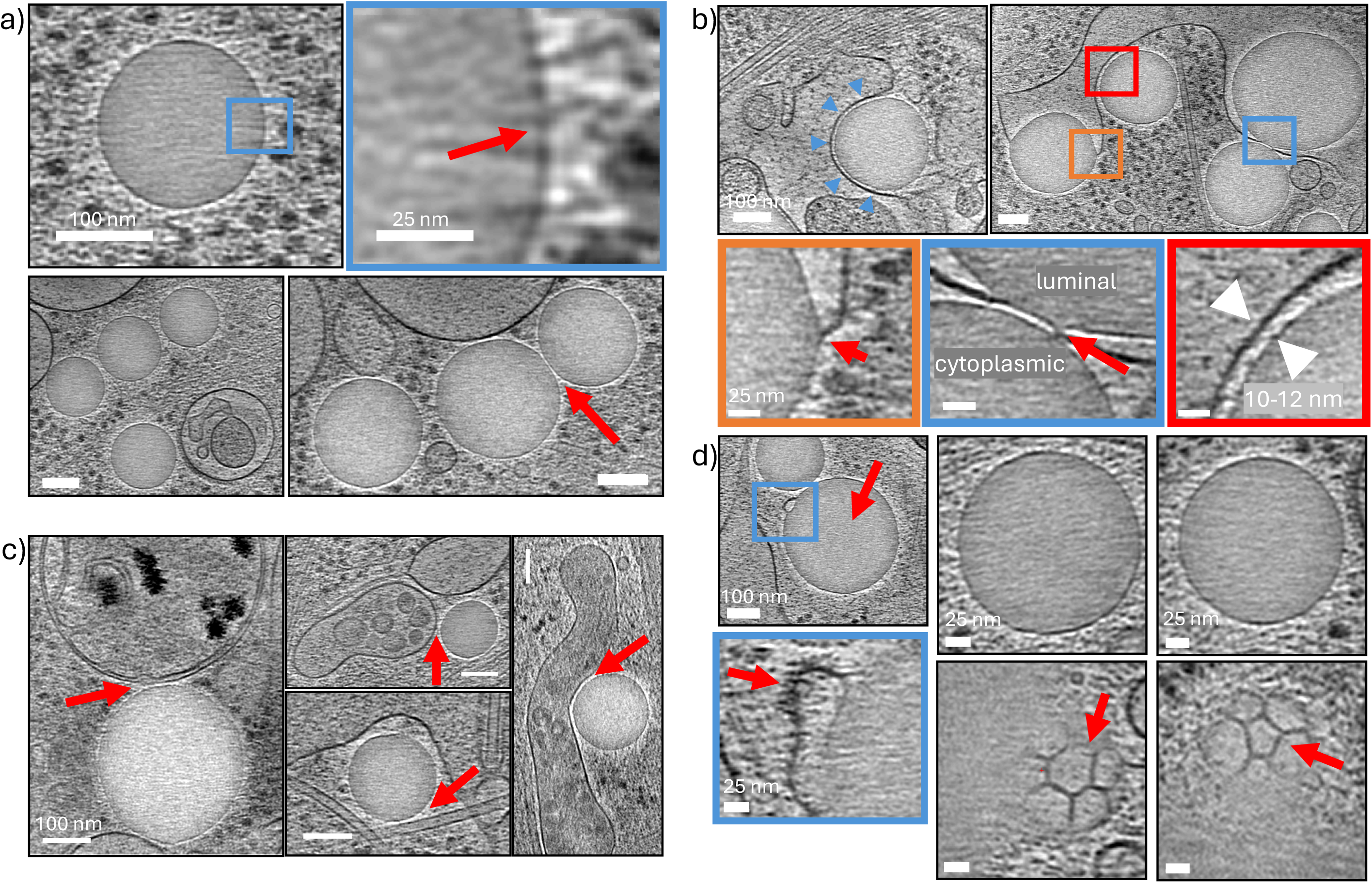
Molecular sociology of lipid droplets (LDs) in U2OS cell periphery as revealed by cryoET at 200 kV. a) LDs were found as spherical, amorphous entities characterized by distinct contrast (top left panel). One of the unique structural features of LDs is their lipid monolayer surface that manifests itself as an electron dense line around perimeter (enlarged view, red arrow). LDs were frequently found near each other (bottom left panel), including direct contact that resulted in flattened and deformed morphology (bottom right panel, red arrow). Scale bars: major panels – 100 nm, enlarged view – 25 nm. b) LDs lifecycle is inherently connected to ER, which is the site of their synthesis. Cytoplasmic LDs were in close association with the ER whose membrane curvature mirrored the LD’s shape (top left panel, blue arrowheads). Three distinct modes of LDs interactions with ER are shown on the top right panel and in enlarged views in the bottom panels, color-coded accordingly. The LD highlighted in orange is embedded within the ER lipid bilayer, which splits to accommodate the bulky volume of the LD (red arrow); in blue are highlighted luminal and cytoplasmic LDs interacting across the ER lipid bilayer (red arrow) and in red a cytoplasmic LD impacting the ER membrane and remaining at a distance of 10-12 nm that is sufficient to accommodate proteins mediating the interaction (Hugenroth & Bohnert, 2020). Scale bars: major panels – 100 nm, enlarged view – 25 nm. c) LDs remain in close association (marked with red arrows) with various cellular modules: mitochondria (left panel), microtubules (middle bottom panel), putative MVBs (middle top panel), membrane embedded compartment that could be a lysosome (right panel). Scale bars: 100 nm. d) LDs interact with membrane trafficking machinery. Top left panel depicts an LD with apparent lipid monolayer deformation (blue square). More detailed analysis (bottom left panel, which is a view of the region marked with the blue square at a different tomographic slice) suggests that the lipid monolayer might be coated with the dense COPI protein layer (red arrow, see also Fig. 2b (II) for comparison). More data and analyses are required to unequivocally identify the protein forming the coat. Top middle and right panels show tomographic slices through central sections of two LDs and below their corresponding views at different tomographic slices. The surface of the two LDs is covered with clathrin that adopts its characteristic architecture with the triskelion forming both hexagonal and pentagonal units (red arrows). Scale bars: top left panel – 100 nm, other panels – 25 nm.

LDs are ubiquitous in the eukaryotic world and essential for lipid and energy metabolism. With the emergence of cryoET imaging we are now beginning to appreciate how LDs interact with other cellular compartments (summarized in **Table 2**), and how these interactions play a role in cellular homeostasis. We believe that structural insights into LDs *in situ* biology obtained by cryoET imaging at 200 kV might further advance the field.

**Table 2.**
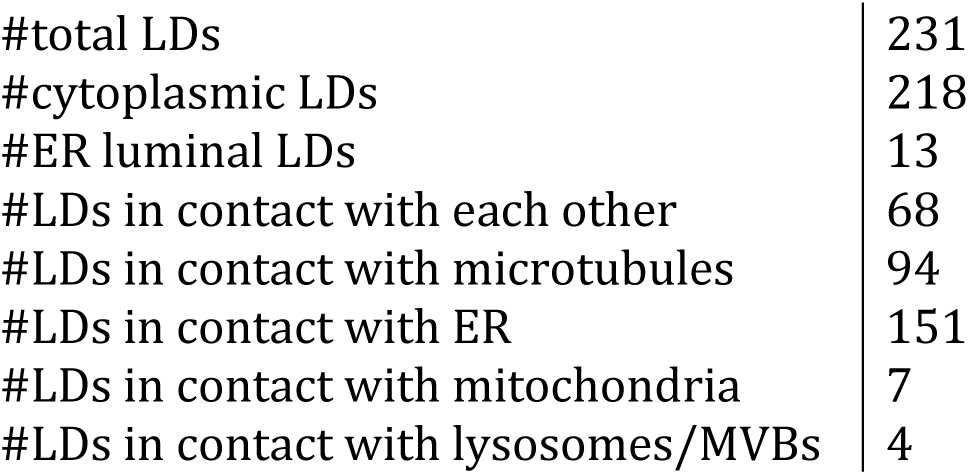
Lipid droplets interactions summary.

## Discussion

CryoET is a versatile and unique structural biology technique as it can bridge information from several orders of magnitude (Å to µm) and is an ideal tool for integrative studies (Ziemianowicz & Kosinski, 2022). In this investigation we sought to evaluate whether a 200 kV cryoTEM equipped with a direct electron detector without an energy filter can deliver interpretable cryoET data that could be used to address structural cell biology problems. Our primary objective was to utilize the mammalian U2OS cell line as the model specimen, compare obtained findings into cellular architecture with the previously published data acquired at 300 kV cryoTEMs fitted with direct electron detectors and energy filters and possibly provide novel insights. It has been demonstrated that radiation damage to biological material is more pronounced at 200 kV than at 300 kV (Peet et al., 2019). Here, we experimentally assessed and carefully tuned the maximal acceptable electron dose to obtain high contrast micrographs suitable for cellular studies by cryoET. This work resulted in several significant outcomes: firstly, the utilized setup allowed for cryoET tilt series acquisition over thin edges of mammalian cells with thicknesses ranging from 50 to 300 nm that could subsequently be reliably reconstructed into 3D volumes; secondly, the resulting tomograms could be reproducibly segmented and all the major cellular components readily recognized; thirdly, the images of RAVs, LDs and other cellular compartments were qualitatively comparable to previously published tomograms acquired by 300 kV cryoTEMs with energy filters (Dudka et al., 2024; Ganeva et al., 2023; Mahamid et al., 2019); fourthly, quantitative analysis of LDs in their native cellular environment revealed the molecular sociology of these organelles. Additionally, in this work we provide molecular resolution direct visual evidence of LDs interaction with membrane trafficking components. We expect that a 200 kV electron cryomicroscope without an energy filter will produce similar results when imaging lamella of cellular/tissue specimens obtained by cryo-focused ion beam milling using either gallium or plasma sources which typically are micromachined to the final thickness of 150-200 nm. It remains to be seen how a 200 kV cryoTEM without an energy filter can perform regarding high-resolution subtomogram averaging, including both *in vitro* and cellular specimens. With this respect we have performed an analysis of purified 70S *Escherichia coli* ribosomes. Subtomogram averaging of **∼**4000 ribosomal particles yielded a 15 Å resolution 3D reconstruction with the threshold of 0.5 for FSC (**Fig. S7**). This demonstrates that a 200 kV cryoEM instrument without an energy filter can generate medium resolution cryoEM maps useful for understanding the overall architecture of macromolecular complexes.

The field of *in situ* structural studies by cryoET is lagging behind SPA cryoEM – as of August 2024 83.4% EMDB entries were derived from SPA analyses vs. 11.4% obtained by cryoET approaches. However, the dynamics of the growth for the cryoET modality is greater – in 2023 year-over-year growth was 50% as compared to 24% for SPA. This rapid growth has been partially possible due to higher throughput data collection strategies (Eisenstein et al., 2023, 2024) and attempts to streamline data processing (Burt et al., 2024; Chen et al., 2019a). Current hardware developments including C_C_ correction, laser phase plates and reduced radiation damage at liquid helium temperatures will further boost the field of cryoET but with a hefty price tag. Such cryoET-purpose built instruments will be prohibitively expensive to purchase and maintain and will be available in a handful of institutions only.

To better understand how current cryoEM resources are utilized we reviewed the output (based on the EMDB database) and distribution (based on the map of high-end cryoEM worldwide (Noble, 2024) – please note that map data might be incomplete since it is based on voluntary contributions from the community) of the two most popular 200 and 300 kV cryoEM platforms: Thermo Scientific Glacios and Thermo Scientific Krios, respectively (**Fig. 4a**). In 2023 there were 5379 EMDB entries attributed to SPA and 837 entries attributed to cryoET obtained from the Krios platform. Whereas Glacios contribution to SPA cryoEM was only 0.95% of Krios in 2020, it steadily rose to 5.9% in 2023. In contrast, the Glacios platform delivered only 4 cryoET EMDB entries in the whole of 2023.

**Figure 4.**
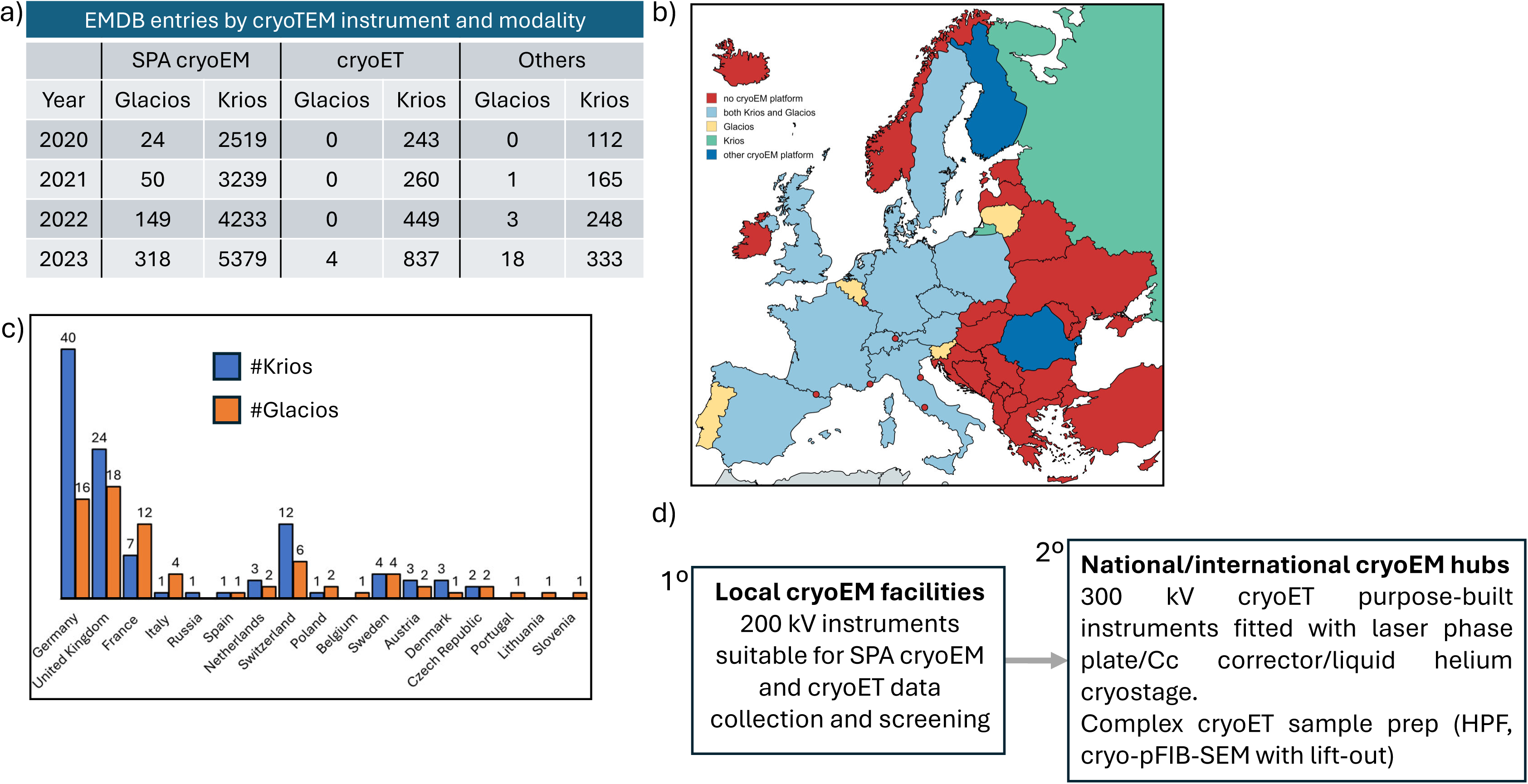
Current utilization of high-end cryoTEMs. a) Table summarizing scientific output generated by two most common 200 kV and 300 kV cryoTEMs: Glacios and Krios, respectively. Whereas the Glacios platform contributed less than 1% of the Krios SPA output in 2020, this fraction rose to 5.9% in 2023. With respect to cryoET, the Glacios platform is heavily underutilized, with only a marginal contribution. Based on the EMDB database as of August 2024. b) Geographical distribution of the Thermo Scientific Glacios and Krios platforms in Europe. As of August 2024, there were ∼102 Krioses and ∼74 Glacioses scattered across 17 European countries. Eastern and South-Eastern Europe regions are underequipped. Please note that in Spain, United Kingdom, Switzerland, Belgium and Germany there are also JEOL cryoARM platform cryoTEMs. Based on the map of high end cryoEM worldwide as of August 2024 (Noble, 2024). c) Number of Glacios and Krios cryoTEMs in European countries. Countries are sorted from left to right according to their nominal gross domestic product (according to IMF, August 2024). d) Implications of this study for core facilities: it is expected that 200 kV instruments will become versatile workhorses in the majority of local cryoEM facilities. These 1^st^ degree access facilities are essential for democratizing access to cryoEM. The 2^nd^ degree access facilities that operate at the national/international level will offer access to cutting-edge imaging technology and advanced sample preparation techniques for cryoET. The technologies for laser phase plates (Schwartz et al., 2019), Cc correctors (J. Wu et al., 2025) and liquid helium cryostages (He et al., 2024) exist but these solutions are not commonly used or not yet available from commercial suppliers.

As of August 2024, there were ∼364 Krioses and ∼230 Glacioses around the world suggesting that 200 kV cryoTEMs are predominantly used as screening devices and heavily underutilized for primary data collection, as evidenced by the number of relevant EMDB entries. In further analysis we focused on Europe that is home to 102 Krioses and 74 Glacioses. As shown on the map, the distribution of these instruments across the continent is not equal. 17 European countries are in possession of either Krios or Glacios or both, 2 countries have access to other cryoEM platforms whereas the 25 remaining countries don’t have access to cryoEM instrumentation at all (**Fig. 4b, c**). The maturity and complexity of cryoEM infrastructure reflects to a certain degree economic development - the top 3 Glacios/Krios owner countries are also 3 top countries by nominal GDP (according to the International Monetary Fund (IMF), August 2024). Next, we surveyed the distribution of the Glacios and Krios cryoEM platforms across academic hubs defined here as cities or agglomerations characterized by the presence of research institutes, either academic or commercial. The Krioses were spread across 39 academic hubs whereas Glacioses across 49, which accounts for 2.6 Krios per hub and 1.5 Glacios per hub. Such clustering of state of the art 300 kV cryoTEMs and better spatial distribution of 200 kV microscopes is related to the fact that the former ones are usually maintained by national/international facilities (eBIC UK, EMBL Germany, NeCEN Netherlands, SciLifeLab Sweden). This trend is visible worldwide, e.g. in the United States there are 3 major SPA cryoEM centers and 4 cryoET centers under the umbrella of NIH Common Fund Transformative High Resolution Cryo-Electron Microscopy Program Centers, each containing multiple high-end instruments. Such centers provide not only access to state-of-the-art microscopes but also to know-how and sample preparation services that, particularly for cellular cryoET, are complex and expensive. The clustering of cryoEM research infrastructure will become more prominent with the arrival of another generation of cryoET purpose-built cryoTEMs. Such microscopes can cost >5 M CHF with added costs for space, service contracts and qualified staff. With this respect it is worth noting that mid-range 200 kV systems have a significantly lower price tag and long-term running costs, e.g. service contracts, which is important in planning facility operations and ensuring optimal instruments performance. The data presented were acquired by operating the Falcon 3EC camera in the linear mode without dose fractionation (due to the impracticalities of long exposure times while operating the Falcon 3EC camera in the counting mode). We suggest that such qualitative studies could also be performed using less costly CMOS-based cameras (e.g. CETA), further bringing down the total investment costs.

200 kV cryoTEMs have been demonstrated to deliver high-quality SPA data, e.g. (Brouwer et al., 2024; Träger et al., 2024) as well as cryoET data from *in vitro* samples, e.g. (Tian et al., 2020). Here, we show that these instruments are capable of delivering excellent cryoET images of cellular specimens. We believe that the 200 kV platform can be a viable option for smaller universities and research institutes that will help democratize cryoEM access and can serve as workhorse for both major cryoEM modalities: SPA and cryoET. We expect that these 1^st^ degree cryoEM access points will further boost the field by attracting top-tier experts, training independent users/operators, facilitating methods development and fostering interdisciplinary research in local academic centers and will greatly complement the 2^nd^ degree cryoEM access points offering the latest and ultimate technological solutions (**Fig. 4d**).

## Acknowledgements

We thank Dr Ishier Raote (CRG Barcelona) for providing the U2OS cell line, Dr Tomasz Góral (University of Warsaw) for the microscope maintenance, Sonia Ngati (University of Warsaw) for the cell culture technical assistance and Dr Fabian Eisenstein (ETH Zurich) for help with setting up and troubleshooting SPACEtomo installation. 70S *E. coli* ribosomes were the generous gift from Dr Alain Scaiola (ETH Zurich). Members of the Center for Microscopy and Image Analysis at the University of Zurich are acknowledged for insightful discussions. This work was supported by the “Regenerative Mechanisms for Health-ReMedy” grant MAB/20172 from the Foundation for Polish Science.

## Author contributions

PS: conception and design, acquisition of data, analysis and interpretation of data, drafting and revising the article

## Supplementary Figures and Movies Legends

### Figures

**Figure S1.**
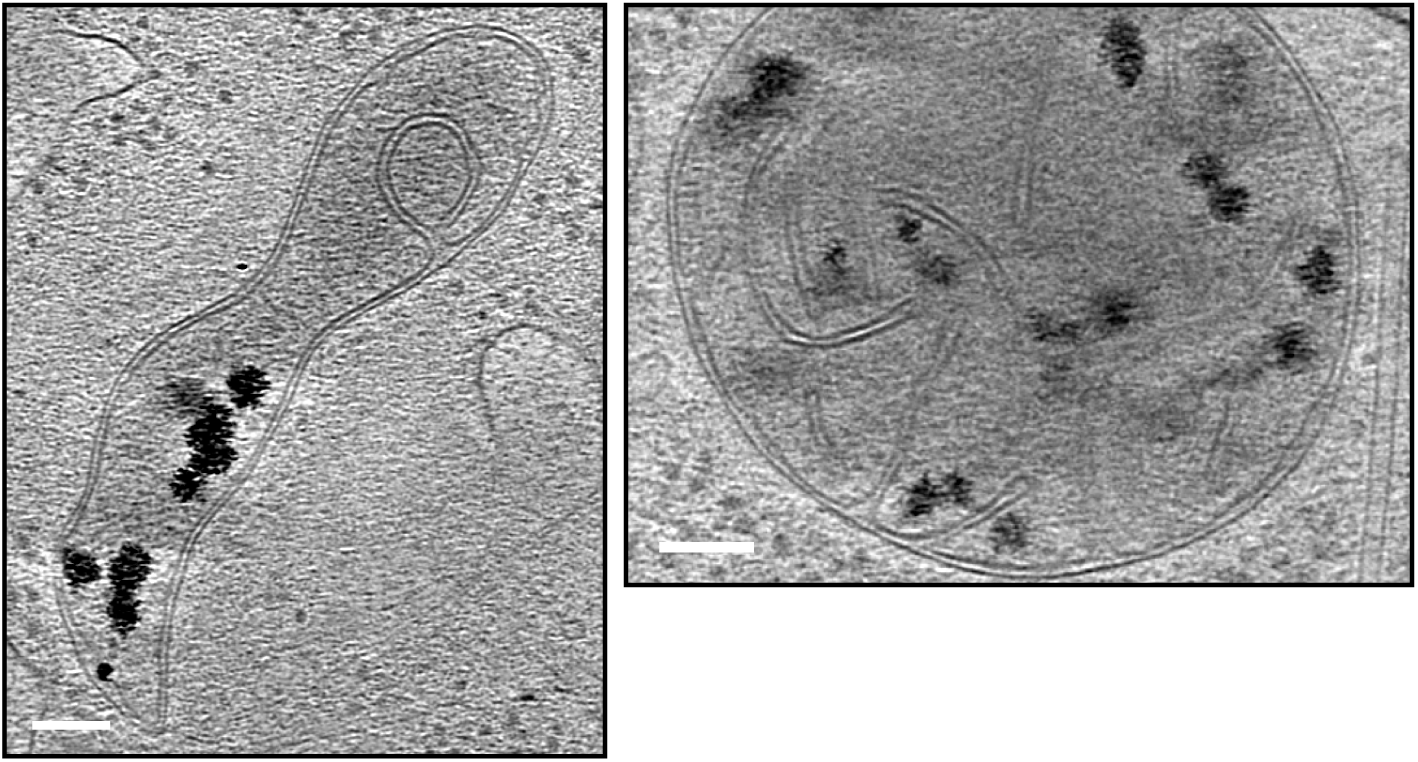
Representative cryoET slices showing mitochondria with calcium phosphate deposits. Corresponding to **Fig. 2b (I)**. Scale bars: 100 nm.

**Figure S2.**
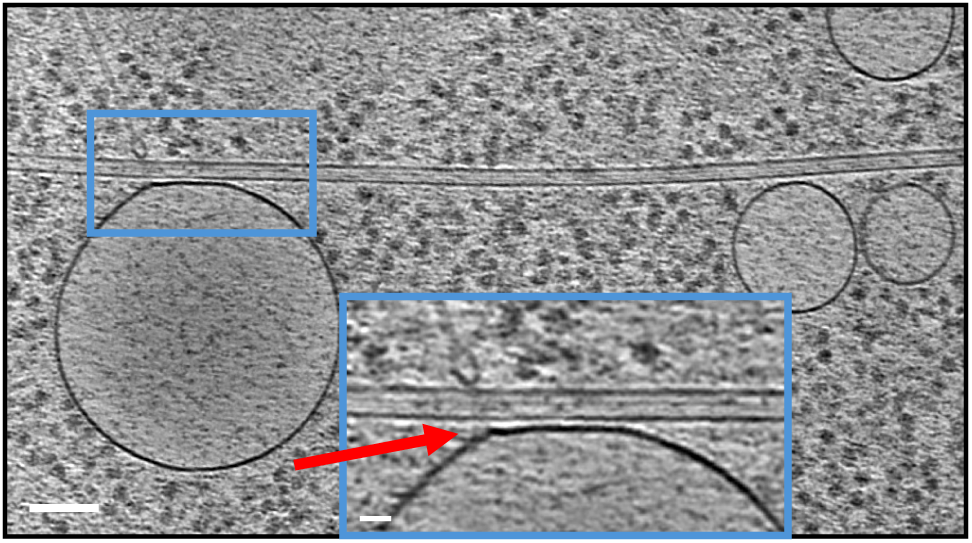
Representative cryoET slice showing a RAV closely associated with a microtubule (red arrow). Corresponding to **Fig. 2b (IV)**. Scale bar: 100 nm.

**Figure S3.**
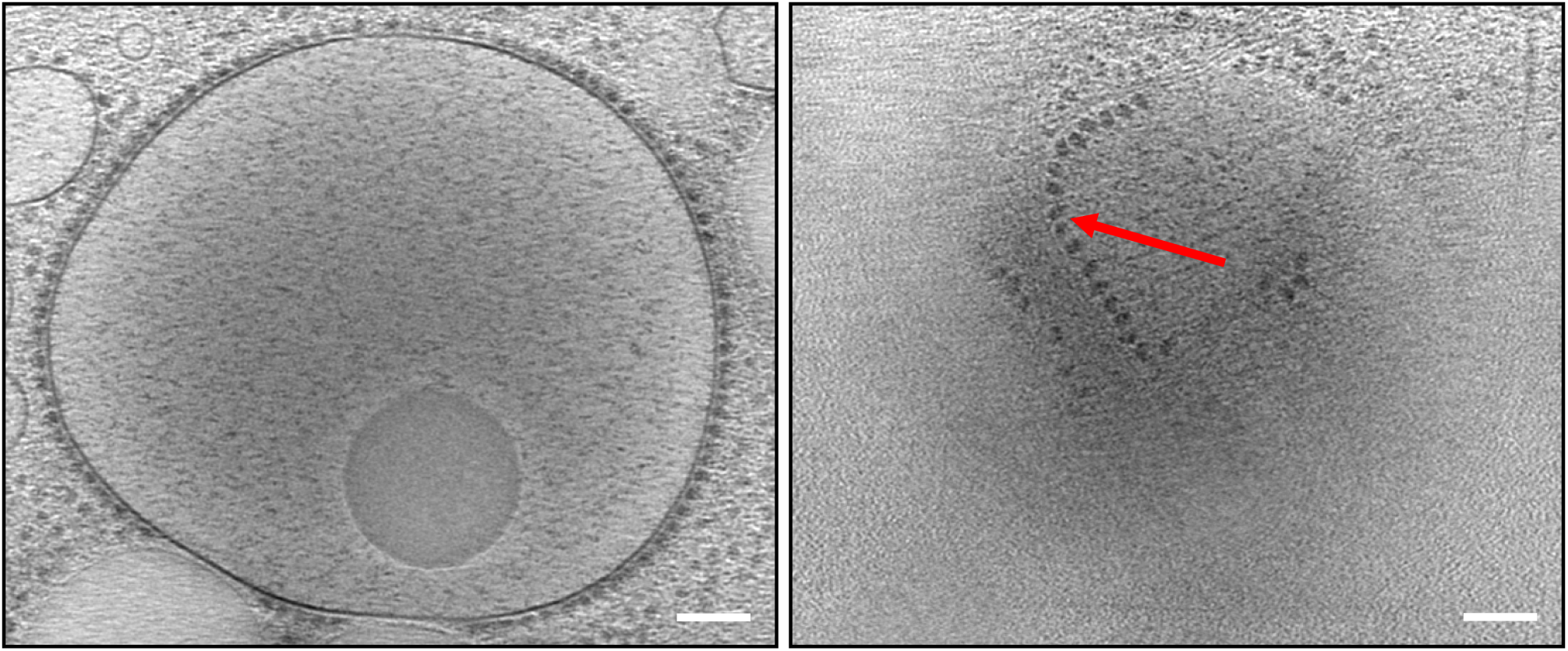
Representative cryoET images showing a RAV at two corresponding views at different tomographic slices. Red arrow highlights a polysome associated with the surface of the RAV. Corresponding to **Fig. 2b (V)** and **Movie S8**. Scale bars: 100 nm.

**Figure S4.**
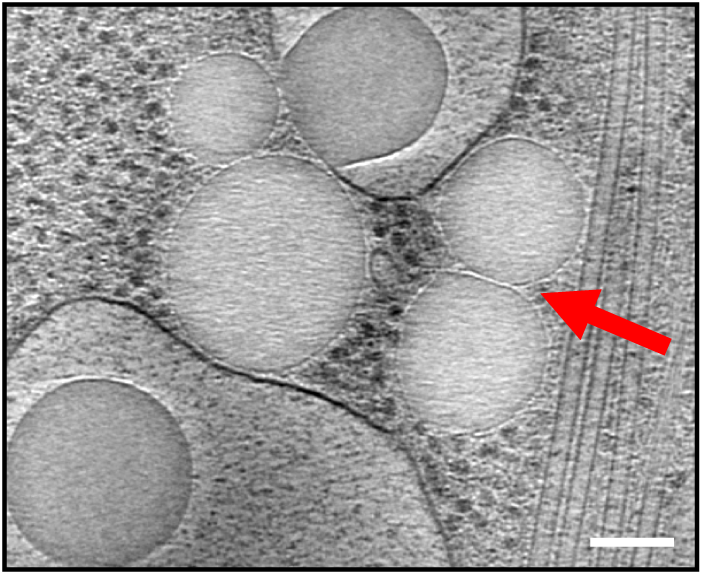
Representative cryoET slice showing several LDs, two of them closely associated with each other (red arrow). Corresponding to **Fig. 3a**. Scale bar: 100 nm.

**Figure S5.**
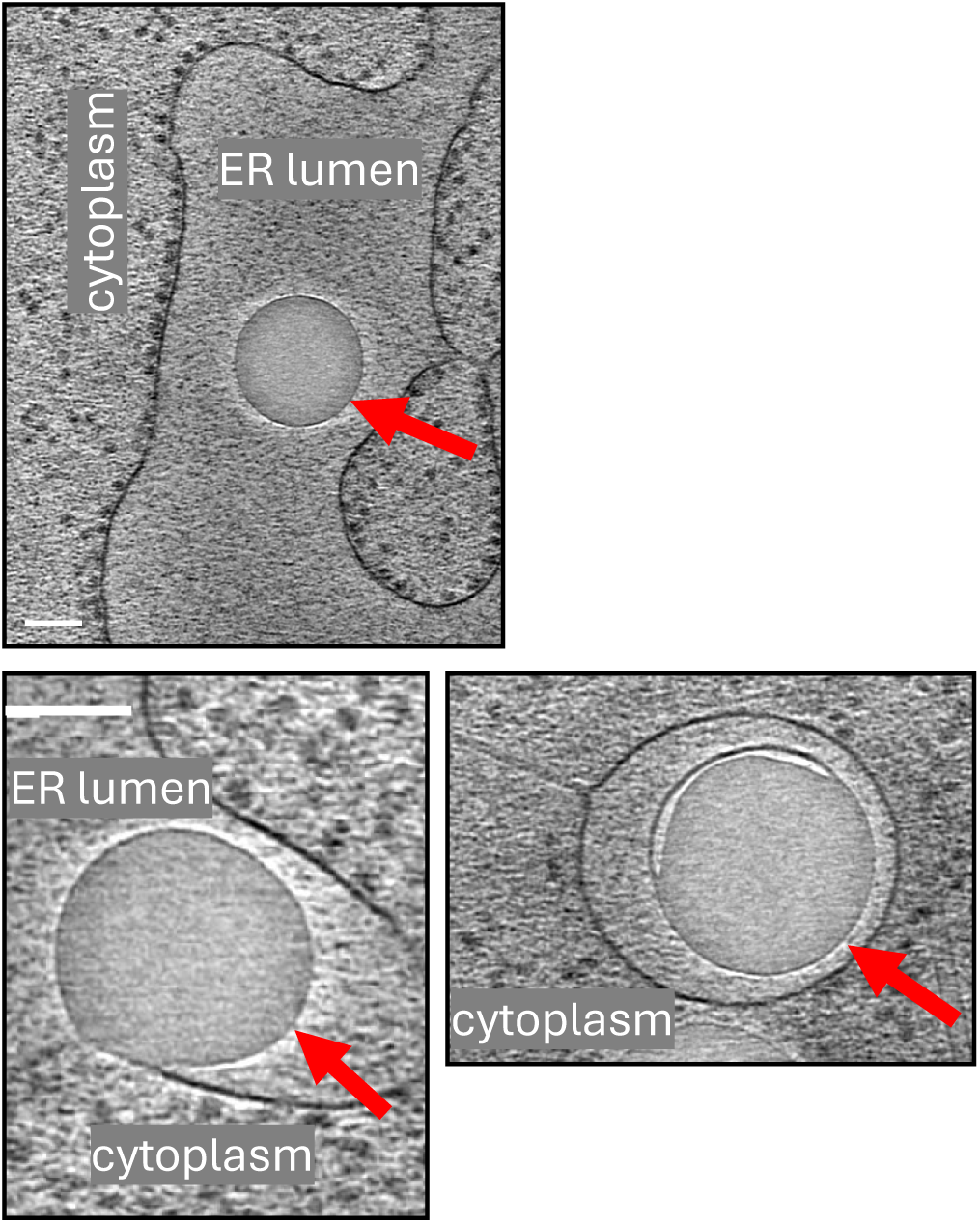
Representative cryoET slices showing LDs residing in the ER lumen. Corresponding to **Fig. 3b**. Scale bars: 100 nm.

**Figure S6.**
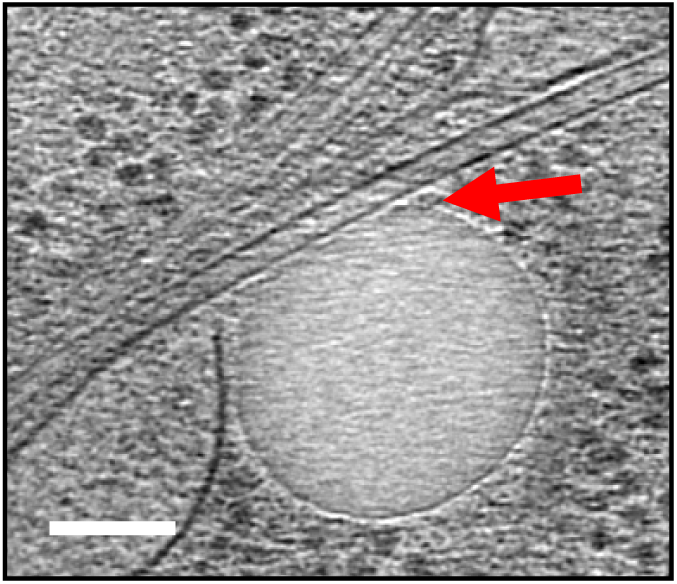
Representative cryoET slice showing LD closely associated with a microtubule (red arrow). Corresponding to **Fig. 3c**. Scale bar: 100 nm.

**Figure S7.**
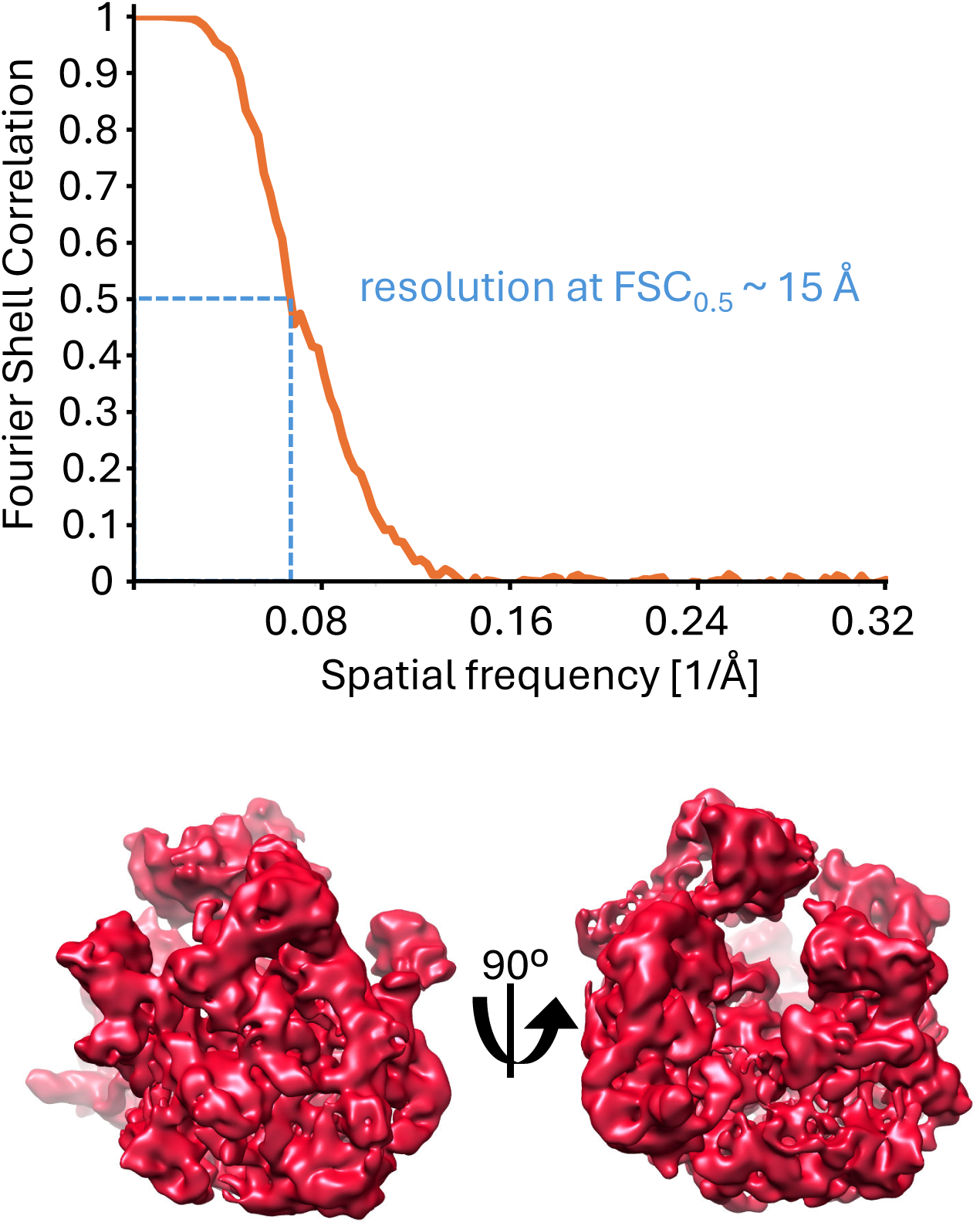
Subtomogram averaging of purified 70S *E. coli* ribosomes. Shown are the FSC curve (top) and overviews of the obtained cryoEM density map.

### Movies

**Movie S1.** Tomogram of a thin edge of an U2OS cell. Corresponding to **Fig. 2a**. Scale bar: 100 nm.

**Movie S2.** Segmentation of a thin edge of an U2OS cell. Corresponding to **Fig. 2a** and **Movie S1**.

**Movie S3.** Tomogram of a mitochondrion found in the thin edge of an U2OS cell. Corresponding to **Fig. 2b (I)**. Scale bar: 100 nm.

**Movie S4.** Tomogram of protein-coated membranes found in the thin edge of an U2OS cell. Corresponding to **Fig. 2b (II)**. Scale bar: 100 nm.

**Movie S5.** Tomogram of ribosome-coated endoplasmic reticulum found in the thin edge of an U2OS cell. Corresponding to **Fig. 2b (III)**. Scale bar: 100 nm.

**Movie S6.** Tomogram of a ribosome-associated vesicle attached to a microtubule found in the thin edge of an U2OS cell. Corresponding to **Fig. 2b (IV)**. Scale bar: 100 nm.

**Movie S7.** Tomogram of two ribosome-associated vesicles connected by a thin membrane tubule found in the thin edge of an U2OS cell. Corresponding to **Fig. 2b (V)**. Scale bar: 100 nm.

**Movie S8.** Tomogram of a large ribosome-associated vesicle with an internal LD and with polysomes positioned on its top surface found in the thin edge of an U2OS cell. Corresponding to **Fig. S3**. Scale bar: 100 nm.

**Movie S9.** Tomogram of a membrane-embedded compartment found in the thin edge of an U2OS cell. Corresponding to **Fig. 2b (VI)**. Scale bar: 100 nm.

**Movie S10.** Tomogram of a membrane-embedded compartment found in the thin edge of an U2OS cell. Corresponding to **Fig. 2b (VII)**. Scale bar: 100 nm.

**Movie S11.** Tomogram of a single lipid droplet found in the thin edge of an U2OS cell. Corresponding to **Fig. 3a**. Scale bar: 100 nm.

**Movie S12.** Tomogram of a bunch of lipid droplets found in the thin edge of an U2OS cell. Corresponding to **Fig. 3a**. Scale bar: 100 nm.

**Movie S13.** Tomogram of lipid droplets closely interacting with an endoplasmic reticulum found in the thin edge of an U2OS cell. Corresponding to **Fig. 3b**. Scale bar: 100 nm.

**Movie S14.** Tomogram of a lipid droplet impacting the lipid bilayer of an endoplasmic reticulum found in the thin edge of an U2OS cell. Corresponding to **Fig. 3b**. Scale bar: 100 nm.

**Movie S15.** Tomogram of a lipid droplet closely interacting with a mitochondrion found in the thin edge of an U2OS cell. Corresponding to **Fig. 3c**. Scale bar: 100 nm.

**Movie S16.** Tomogram of a lipid droplet closely interacting with microtubules found in the thin edge of an U2OS cell. Corresponding to **Fig. 3c**. Scale bar: 100 nm.

**Movie S17.** Tomogram of a lipid droplet putatively coated with COPI found in the thin edge of an U2OS cell. Corresponding to **Fig. 3d**. Scale bar: 25 nm.

**Movie S18 and Movie S19.** Tomograms of lipid droplets coated with clathrin found in the thin edge of an U2OS cell. Corresponding to **Fig. 3d**. Scale bars: 25 nm.

## Methods

### Sample preparation

Samples for cellular electron cryotomography were prepared as follows: Quantifoil Au200 R2/2 grids were glow discharged for 90 s at 20 mA, coated with fibronectin (25 μg/ml) and transferred to a 12 well plate in 3D printed cell culture grid holders to facilitate handling (Fäßler et al., 2020). After sterilization for 30 min under UV light, the grids were seeded with U2OS cells (∼6000 per well) and incubated at 37⁰C overnight. Before plunge freezing the grids were briefly washed in PBS and 2.5 μl of 10 nm colloidal gold fiducial markers was applied. Subsequently, the grids were back-blotted and vitrified in liquid ethane using a Thermo Scientific Vitrobot.

Samples for electron cryotomography of purified macromolecular complexes were prepared as follows: Quantifoil Au200 R1.2/1.3 grids were glow discharged for 90 s at 20 mA. Then 4 μl of *Escherichia coli* 70S ribosomes at 250 nM was applied and vitrified in liquid ethane-propane using a Leica EM GP2.

### Data collection

Cellular electron cryotomography was performed using a Thermo Scientific Glacios G1 TEM operating at 200 kV, equipped with a 4k x 4k Falcon 3EC direct electron detection camera set in the linear mode without dose fractionation at a magnification of 28k with parallel illumination conditions, corresponding to a pixel size of 5.1 Å at the specimen level. Samples were tilted between approximately −54⁰ and +54⁰ with a 3⁰ increment using the dose symmetric scheme in the TOMO software. The defocus was set between −5 and −7 μm and the total dose for each tilt series was 140 eÅ^-2^.

Electron cryotomography of the 70S ribosomes was performed using a Thermo Scientific Glacios G2 TEM operating at 200 kV, equipped with a 4k x 4k Falcon 4i direct electron detection camera set in the counting mode at a magnification of 92k with parallel illumination conditions, corresponding to a pixel size of 1.6 Å at the specimen level. Samples were tilted between approximately −54⁰ and +54⁰ with a 3⁰ increment using the dose symmetric scheme and beam image shift data collection strategy implemented in the (S)PACEtomo/SerialEM software was used to record tilt series (Eisenstein et al., 2023, 2024; Mastronarde, 2005). The defocus was set between −1.5 and −3 μm and 1.25 s exposures were divided into 8 frames each. The total dose for each tilt series was 130 eÅ^-2^.

### Data analysis

Tomographic reconstruction from tilt series were calculated using RAPTOR (Amat et al., 2008) and the IMOD tomography reconstruction package (Kremer et al., 1996) followed by Weighted Back Projection or Simultaneous Iterative Reconstruction Technique (SIRT) with the tomo3d package (Agulleiro & Fernandez, 2011). Measurements of distances, including tomogram thicknesses, were carried out within IMOD. Here, two model points were placed (to mark the top and bottom surface of the biological density in the tomogram) and the distance in 3D between them calculated in 3dmod. Single threshold surface representation and movies were prepared using Chimera, with a similar threshold level to exclude most background noise for each data set. EMAN2 (Chen et al., 2017, 2019b) was used for tomogram segmentations/annotations with the use of the neural network and subtomogram averaging of *in situ* and purified ribosomes. The neural network was trained to recognize different cellular features by selecting 5 tomograms with slightly various defocus values, then identifying in every tomogram ∼20 tiles containing the feature of interest and ∼100 tiles deprived of the feature, after which the content of the tiles was manually annotated and fed into the neural network. CTFFIND4 (Rohou & Grigorieff, 2015) was used to obtain information about CTF fit.

### Data deposition

Representative 3D volumes have been deposited in the Electron Microscopy Data Bank (EMDB) under accession numbers: EMD-51957, EMD-51959, EMD-51960, EMD-51961, EMD-51962 and EMD-54370 and representative tilt series in the EMPIAR database under accession numbers: EMPIAR-12763, EMPIAR-12764, EMPIAR-12765, EMPIAR-12766 and EMPIAR-12767.

## Declarations

### Ethics approval and consent to participate

Not applicable, as no patient data was used in this research. Cell line used in this study is not relevant material under the Human Research Act, so no ethical approval was required.

### Consent for publication

Not applicable

### Availability of data and material

Data is provided within the manuscript or supplementary information files. Additionally, representative 3D volumes have been deposited in the Electron Microscopy Data Bank (EMDB) under accession numbers: EMD-51957, EMD-51959, EMD-51960, EMD-51961, EMD-51962 and EMD-54370 and representative tilt series in the EMPIAR database under accession numbers: EMPIAR-12763, EMPIAR-12764, EMPIAR-12765, EMPIAR-12766 and EMPIAR-12767.

## References

Agulleiro, J. I., & Fernandez, J. J. (2011). Fast tomographic reconstruction on multicore computers. Bioinformatics, 27(4), 582–583. 10.1093/bioinformatics/btq692

Amat, F., Moussavi, F., Comolli, L. R., Elidan, G., Downing, K. H., & Horowitz, M. (2008). Markov random field based automatic image alignment for electron tomography. Journal of Structural Biology, 161(3), 260–275. 10.1016/j.jsb.2007.07.007

Bartz, R., Zehmer, J. K., Zhu, M., Chen, Y., Serrero, G., Zhao, Y., & Liu, P. (2007). Dynamic activity of lipid droplets: Protein phosphorylation and GTP-mediated protein translocation. Journal of Proteome Research, 6(8), 3256– 3265. 10.1021/pr070158j

Beck, M., & Baumeister, W. (2016). Cryo-Electron Tomography: Can it Reveal the Molecular Sociology of Cells in Atomic Detail? In Trends in Cell Biology (Vol. 26, Issue 11, pp. 825–837). Elsevier Ltd. 10.1016/j.tcb.2016.08.006

Brouwer, P. J. M., Perrett, H. R., Beaumont, T., Nijhuis, H., Kruijer, S., Burger, J. A., Bontjer, I., Lee, W.-H., Ferguson, J. A., Schauflinger, M., Müller-Kräuter, H., Sanders, R. W., Strecker, T., van Gils, M. J., & Ward, A. B. (2024). Defining bottlenecks and opportunities for Lassa virus neutralization by structural profiling of vaccine-induced polyclonal antibody responses. Cell Reports, 43(9), 114708. 10.1016/j.celrep.2024.114708

Burt, A., Toader, B., Warshamanage, R., von Kügelgen, A., Pyle, E., Zivanov, J., Kimanius, D., Bharat, T. A. M., & Scheres, S. H. W. (2024). An image processing pipeline for electron cryo-tomography in RELION-5. FEBS Open Bio. 10.1002/2211-5463.13873

Bykov, Y. S., Schaffer, M., Dodonova, S. O., Albert, S., rgen Plitzko, J. M., Baumeister, W., Engel, B. D., & Briggs, J. A. (2017). The structure of the COPI coat determined within the cell. 10.7554/eLife.32493.001

Carter, S. D., Hampton, C. M., Langlois, R., Melero, R., Farino, Z. J., Calderon, M. J., Li, W., Wallace, C. T., Han Tran, N., Grassucci, R. A., Siegmund, S. E., Pemberton, J., Morgenstern, T. J., Bartolini, F., Volchuk, A., Murray, S. A., Aridor, M., Fish, K. N., Walter, P., … Freyberg, Z. (2020). Ribosome-associated vesicles: A dynamic subcompartment of the endoplasmic reticulum in secretory cells. In Despoina Aslanoglou (Vol. 10). https://www.science.org

Chan, L. M., Courteau, B. J., Maker, A., Wu, M., Basanta, B., Mehmood, H., Bulkley, D., Joyce, D., Lee, B. C., Mick, S., Czarnik, C., Gulati, S., Lander, G. C., & Verba, K. A. (2024). High-resolution single-particle imaging at 100–200 keV with the Gatan Alpine direct electron detector. Journal of Structural Biology, 216(3). 10.1016/j.jsb.2024.108108

Chen, M., Bell, J. M., Shi, X., Sun, S. Y., Wang, Z., & Ludtke, S. J. (2019a). A complete data processing workflow for cryo-ET and subtomogram averaging. Nature Methods, 16(11), 1161–1168. 10.1038/s41592-019-0591-8

Chen, M., Bell, J. M., Shi, X., Sun, S. Y., Wang, Z., & Ludtke, S. J. (2019b). A complete data processing workflow for cryo-ET and subtomogram averaging. Nature Methods, 16(11), 1161–1168. 10.1038/s41592-019-0591-8

Chen, M., Dai, W., Sun, S. Y., Jonasch, D., He, C. Y., Schmid, M. F., Chiu, W., & Ludtke, S. J. (2017). Convolutional neural networks for automated annotation of cellular cryo-electron tomograms. Nature Methods, 14(10), 983–985. 10.1038/nmeth.4405

Daro, E., Sheff, D., Gomez, M., Kreis, T., & Mellman, I. (1997). Inhibition of Endosome Function in CHO Cells Bearing a Temperature-sensitive Defect in the Coatomer (COPI) Component-COP. In The Journal of Cell Biology (Vol. 139, Issue 7). http://www.jcb.org

Dudka, W., Salo, V. T., & Mahamid, J. (2024). Zooming into lipid droplet biology through the lens of electron microscopy. In FEBS Letters (Vol. 598, Issue 10, pp. 1127–1142). John Wiley and Sons Inc. 10.1002/1873-3468.14899

Eisenstein, F., Fukuda, Y., & Danev, R. (2024). Smart parallel automated cryo-electron tomography. Nature Methods, 21(9), 1612–1615. 10.1038/s41592-024-02373-9

Eisenstein, F., Yanagisawa, H., Kashihara, H., Kikkawa, M., Tsukita, S., & Danev, R. (2023). Parallel cryo electron tomography on in situ lamellae. Nature Methods, 20(1), 131–138. 10.1038/s41592-022-01690-1

Faini, M., Prinz, S., Beck, R., Schorb, M., Riches, J. D., Bacia, K., Brügger, B., Wieland, F. T., & Briggs, J. A. G. (2012). The structures of COPI-coated vesicles reveal alternate coatomer conformations and interactions. Science, 336(6087), 1451–1454. 10.1126/science.1221443

Fan, H., & Tan, Y. (2024). Lipid Droplet–Mitochondria Contacts in Health and Disease. In International Journal of Molecular Sciences (Vol. 25, Issue 13). Multidisciplinary Digital Publishing Institute (MDPI). 10.3390/ijms25136878

Fäßler, F., Zens, B., Hauschild, R., & Schur, F. K. M. (2020). 3D printed cell culture grid holders for improved cellular specimen preparation in cryo-electron microscopy. Journal of Structural Biology, 212(3). 10.1016/j.jsb.2020.107633

Förster, F., Han, B. G., & Beck, M. (2010). Visual Proteomics. In Methods in Enzymology (Vol. 483, Issue C, pp. 215–243). Academic Press Inc. 10.1016/S0076-6879(10)83011-3

Foster, H. E., Santos, C. V., & Carter, A. P. (2022). A cryo-ET survey of microtubules and intracellular compartments in mammalian axons. Journal of Cell Biology, 221(2). 10.1083/jcb.202103154

Fotin, A., Kirchhausen, T., Grigorieff, N., Harrison, S. C., Walz, T., & Cheng, Y. (2006). Structure determination of clathrin coats to subnanometer resolution by single particle cryo-electron microscopy. Journal of Structural Biology, 156(3), 453–460. 10.1016/j.jsb.2006.07.001

Fry, M. Y., Navarro, P. P., Hakim, P., Ananda, V. Y., Qin, X., Landoni, J. C., Rath, S., Inde, Z., Lugo, C. M., Luce, B. E., Ge, Y., Mcdonald, J. L., Ali, I., Ha, L. L., Kleinstiver, B. P., Chan, D. C., Sarosiek, K. A., & Chao, L. H. (2024). In situ architecture of Opa1-dependent mitochondrial cristae remodeling. 10.1101/2023.01.16.524176

Fujimoto, T., & Parton, R. G. (2011). Not just fat: The structure and function of the lipid droplet. Cold Spring Harbor Perspectives in Biology, 3(3), 1–17. 10.1101/cshperspect.a004838

Ganeva, I., Lim, K., Boulanger, J., Hoffmann, P. C., Muriel, O., Borgeaud, A. C., Hagen, W. J. H., Savage, D. B., & Kukulski, W. (2023). The architecture of Cidec-mediated interfaces between lipid droplets. Cell Reports, 42(2). 10.1016/j.celrep.2023.112107

Gross, S. P., Welte, M. A., Block, S. M., & Wieschaus, E. F. (2000). Dynein-mediated Cargo Transport In Vivo: A Switch Controls Travel Distance. In The Journal of Cell Biology (Vol. 148, Issue 5). http://www.jcb.org

He, X., Kostin, R., Knight, E., Han, M. G., Mun, J., Bozovic, I., Jing, C., & Zhu, Y. (2024). Development of a liquid-helium free cryogenic sample holder with mK temperature control for autonomous electron microscopy. Ultramicroscopy, 267. 10.1016/j.ultramic.2024.114037

Herzik, M. A., Wu, M., & Lander, G. C. (2017). Achieving better-than-3-Å resolution by single-particle cryo-EM at 200 keV. Nature Methods, 14(11), 1075–1078. 10.1038/nmeth.4461

Hugenroth, M., & Bohnert, M. (2020). Come a little bit closer! Lipid droplet-ER contact sites are getting crowded. In Biochimica et Biophysica Acta - Molecular Cell Research (Vol. 1867, Issue 2). Elsevier B.V. 10.1016/j.bbamcr.2019.118603

Kilwein, M. D., & Welte, M. A. (2019). Lipid droplet motility and organelle contacts. Contact (Thousand Oaks), 585, 276–296.

Kremer, J. R., Mastronarde, D. N., & Mcintosh, J. R. (1996). Computer Visualization of Three-Dimensional Image Data Using IMOD.

Kühlbrandt, W. (2014). The resolution revolution. In Science (Vol. 343, Issue 6178, pp. 1443–1444). American Association for the Advancement of Science. 10.1126/science.1251652

Mahamid, J., Tegunov, D., Maiser, A., Arnold, J., Leonhardt, H., Plitzko, J. M., & Baumeister, W. (2019). Liquid-crystalline phase transitions in lipid droplets are related to cellular states and specific organelle association. Proceedings of the National Academy of Sciences of the United States of America, 116(34), 16866–16871. 10.1073/pnas.1903642116

Martynowycz, M. W., Clabbers, M. T. B., Unge, J., Hattne, J., & Gonen, T. (2021). Benchmarking the ideal sample thickness in cryo-EM. 118, 2108884118. 10.1073/pnas.2108884118/-/DCSupplemental

Mastronarde, D. N. (2005). Automated electron microscope tomography using robust prediction of specimen movements. Journal of Structural Biology, 152(1), 36–51. 10.1016/j.jsb.2005.07.007

McMullan, G. I., Naydenova, K. I., Mihaylov, D., Yamashita, K. I., Peet ID, M. J., Wilson, H. I., Dickerson ID, J. L., Chen, S. I., Cannone, G. I., Lee, Y. I., Hutchings ID, K. A., Gittins, O. I., Sobhy ID, M. A., Wells, T. I., El-Gomati ID, M. M., Dalby, J., Meffert, M. I., Schulze-Briese, C. I., Henderson, R. I., & Russo, C. J. (2023). Structure determination by cryoEM at 100 keV. 10.1073/pnas

Medalia, O., Weber, I., Frangakis, A. S., Nicastro, D., Gerisch, G., & Baumeister, W. (2002). Macromolecular Architecture in Eukaryotic Cells Visualized by Cryoelectron Tomography. Science, 5596, 1209–1213. DOI:10.1126/science.1076184

Navaratnarajah, T., Anand, R., Reichert, A. S., & Distelmaier, F. (2021). The relevance of mitochondrial morphology for human disease. In International Journal of Biochemistry and Cell Biology (Vol. 134). Elsevier Ltd. 10.1016/j.biocel.2021.105951

Noble, A. J. (2024). HighEnd CryoEM Worldwide.

Nogales, E., & Mahamid, J. (2024). Bridging structural and cell biology with cryo-electron microscopy. In Nature (Vol. 628, Issue 8006, pp. 47–56). Nature Research. 10.1038/s41586-024-07198-2

Nonoyama, T., Nojima, D., Maeda, Y., Noda, M., Yoshino, T., Matsumoto, M., Bowler, C., & Tanaka, T. (2019). Proteomics analysis of lipid droplets indicates involvement of membrane trafficking proteins in lipid droplet breakdown in the oleaginous diatom Fistulifera solaris. Algal Research, 44. 10.1016/j.algal.2019.101660

Olarte, M. J., Swanson, J. M. J., Walther, T. C., & Farese, R. V. (2022). The CYTOLD and ERTOLD pathways for lipid droplet–protein targeting. In Trends in Biochemical Sciences (Vol. 47, Issue 1, pp. 39–51). Elsevier Ltd. 10.1016/j.tibs.2021.08.007

Olzmann, J. A., & Carvalho, P. (2019). Dynamics and functions of lipid droplets. In Nature Reviews Molecular Cell Biology (Vol. 20, Issue 3, pp. 137–155). Nature Publishing Group. 10.1038/s41580-018-0085-z

Peet, M. J., Henderson, R., & Russo, C. J. (2019). The energy dependence of contrast and damage in electron cryomicroscopy of biological molecules. Ultramicroscopy, 203, 125–131. 10.1016/j.ultramic.2019.02.007

Punjani, A., & Fleet, D. J. (2023). 3DFlex: determining structure and motion of flexible proteins from cryo-EM. Nature Methods, 20(6), 860–870. 10.1038/s41592-023-01853-8

Rice, W. J., Cheng, A., Noble, A. J., Eng, E. T., Kim, L. Y., Carragher, B., & Potter, C. S. (2018). Routine determination of ice thickness for cryo-EM grids. Journal of Structural Biology, 204(1), 38–44. 10.1016/j.jsb.2018.06.007

Rigort, A., & Plitzko, J. M. (2015). Cryo-focused-ion-beam applications in structural biology. In Archives of Biochemistry and Biophysics (Vol. 581, pp. 122–130). Academic Press Inc. 10.1016/j.abb.2015.02.009

Rohou, A., & Grigorieff, N. (2015). CTFFIND4: Fast and accurate defocus estimation from electron micrographs. Journal of Structural Biology, 192(2), 216–221. 10.1016/j.jsb.2015.08.008

Russo, C. J., Dickerson, J. L., & Naydenova, K. (2022). Cryomicroscopy in situ: what is the smallest molecule that can be directly identified without labels in a cell? Faraday Discussions, 240, 277–302. 10.1039/d2fd00076h

Schwab, J., Kimanius, D., Burt, A., Dendooven, T., & Scheres, S. H. W. (2024). DynaMight: estimating molecular motions with improved reconstruction from cryo-EM images. Nature Methods. 10.1038/s41592-024-02377-5

Schwartz, O., Axelrod, J. J., Campbell, S. L., Turnbaugh, C., Glaeser, R. M., & Müller, H. (2019). Laser phase plate for transmission electron microscopy. Nature Methods, 16(10), 1016–1020. 10.1038/s41592-019-0552-2

Szwedziak, P., & Pilhofer, M. (2019). Bidirectional contraction of a type six secretion system. Nature Communications, 10(1). 10.1038/s41467-019-09603-1

Szwedziak, P., Wang, Q., Bharat, T. A., Tsim, M., & Löwe, J. (2014). Architecture of the ring formed by the tubulin homologue FtsZ in bacterial cell division. ELife, 3. 10.7554/eLife.04601

Tegunov, D., Xue, L., Dienemann, C., Cramer, P., & Mahamid, J. (2021). Multi-particle cryo-EM refinement with M visualizes ribosome-antibiotic complex at 3.5 Å in cells. Nature Methods, 18(2), 186–193. 10.1038/s41592-020-01054-7

Thiam, A. R., Antonny, B., Wang, J., Delacotte, J., Wilfling, F., Walther, T. C., Beck, R., Rothman, J. E., & Pincet, F. (2013). COPI buds 60-nm lipid droplets from reconstituted water-phospholipid-triacylglyceride interfaces, suggesting a tension clamp function. Proceedings of the National Academy of Sciences of the United States of America, 110(33), 13244–13249. 10.1073/pnas.1307685110

Tian, Y., Liang, R., Kumar, A., Szwedziak, P., & Viles, J. H. (2020). 3D-visualization of amyloid-β oligomer and fibril interactions with lipid membranes by cryo-electron tomography. In bioRxiv. 10.1101/2020.07.21.214072

Träger, T. K., Kyrilis, F. L., Hamdi, F., Tüting, C., Alfes, M., Hofmann, T., Schmidt, C., & Kastritis, P. L. (2024). Disorder-to-order active site capping regulates the rate-limiting step of the inositol pathway. Proceedings of the National Academy of Sciences of the United States of America, 121(34). 10.1073/pnas.2400912121

Valm, A. M., Cohen, S., Legant, W. R., Melunis, J., Hershberg, U., Wait, E., Cohen, A. R., Davidson, M. W., Betzig, E., & Lippincott-Schwartz, J. (2017). Applying systems-level spectral imaging and analysis to reveal the organelle interactome. In Nature (Vol. 546, Issue 7656, pp. 162–167). Nature Publishing Group. 10.1038/nature22369

Wilfling, F., Thiam, A. R., Olarte, M. J., Wang, J., Beck, R., Gould, T. J., Allgeyer, E. S., Pincet, F., Bewersdorf, J., Farese, R. V., & Walther, T. C. (2014). Arf1/COPI machinery acts directly on lipid droplets and enables their connection to the ER for protein targeting. ELife, 2014(3). 10.7554/eLife.01607

Wilfling, F., Wang, H., Haas, J. T., Krahmer, N., Gould, T. J., Uchida, A., Cheng, J. X., Graham, M., Christiano, R., Fröhlich, F., Liu, X., Buhman, K. K., Coleman, R. A., Bewersdorf, J., Farese, R. V., & Walther, T. C. (2013). Triacylglycerol synthesis enzymes mediate lipid droplet growth by relocalizing from the ER to lipid droplets. Developmental Cell, 24(4), 384–399. 10.1016/j.devcel.2013.01.013

Wolf, S., Mutsafi, yael, Dadosh, T., Ilani, T., Lansky, Z., Horowitz, B., Rubin, S., Elbaum, M., & Fass, D. (2017). 3D visualization of mitochondrial solid-phase calcium stores in whole cells. ELife. 10.7554/eLife.29929.001

Wolff, G., Limpens, R. W. A. L., Zevenhoven-Dobbe, J. C., Laugks, U., Zheng, S., De Jong, A. W. M., Koning, R. I., Agard, D. A., Grünewald, K., Koster, A. J., Snijder, E. J., & Bárcena, M. (2020). A molecular pore spans the double membrane of the coronavirus replication organelle. Science, 1395–1398. 10.1126/science.abd3629

Wozny, M. R., Di Luca, A., Morado, D. R., Picco, A., Khaddaj, R., Campomanes, P., Ivanović, L., Hoffmann, P. C., Miller, E. A., Vanni, S., & Kukulski, W. (2023). In situ architecture of the ER–mitochondria encounter structure. Nature, 618(7963), 188–192. 10.1038/s41586-023-06050-3

Wu, G. H., Smith-Geater, C., Galaz-Montoya, J. G., Gu, Y., Gupte, S. R., Aviner, R., Mitchell, P. G., Hsu, J., Miramontes, R., Wang, K. Q., Geller, N. R., Hou, C., Danita, C., Joubert, L. M., Schmid, M. F., Yeung, S., Frydman, J., Mobley, W., Wu, C., … Chiu, W. (2023). CryoET reveals organelle phenotypes in huntington disease patient iPSC-derived and mouse primary neurons. Nature Communications, 14(1). 10.1038/s41467-023-36096-w

Wu, J., Liu, C., Wang, A. J., Gao, Y. Z., Fu, L. T., Liu, Z., Dickerson, J. L., Russo, C. J., & Wang, P. (2025). Chromatic aberration (Cc) corrected cryo-EM: The structure of pseudorabies virus (PRV) using both zero-loss and energy loss electrons. Ultramicroscopy, 276. 10.1016/j.ultramic.2025.114182

Zhang, S., Peng, X., Yang, S., Li, X., Huang, M., Wei, S., Liu, J., He, G., Zheng, H., Yang, L., Li, H., & Fan, Q. (2022). The regulation, function, and role of lipophagy, a form of selective autophagy, in metabolic disorders. In Cell Death and Disease (Vol. 13, Issue 2). Springer Nature. 10.1038/s41419-022-04593-3

Ziemianowicz, D. S., & Kosinski, J. (2022). New opportunities in integrative structural modeling. In Current Opinion in Structural Biology (Vol. 77). Elsevier Ltd. 10.1016/j.sbi.2022.102488

